# Adrenal sympathetic nerve mediated the anti-inflammatory effect of electroacupuncture at ST25 acupoint on a rat model of sepsis

**DOI:** 10.1101/2022.07.14.499985

**Authors:** Ziyi Zhang, Xiang Cui, Kun Liu, Xinyan Gao, QingChen Zhou, Hanqing Xi, Yingkun Zhao, Dingdan Zhang, Bing Zhu

**Affiliations:** Department of Physiology, Institute of Acupuncture and Moxibustion, China Academy of Chinese Medical Sciences, Beijing, 100700, China; College of Acupuncture and Tuina, Shaanxi University of Chinese Medicine, Xianyang, Shaanxi Province, 712046, China; Acupuncture-moxibustion and Tuina Department, Qilu Hospital of Shandong University, Jinan, Shandong, 250014, China; Third Affiliated Hospital, Beijing University of Chinese Medicine, Beijing, 100039, China; Dongzhimen Hospital, Beijing University of Chinese Medicine, Beijing, 100700, China

**Keywords:** Autonomic nerve, Neuro-immune, Noradrenaline, Neuromodulation, Sympathoadrenal medullary pathway

## Abstract

Acupuncture plays a vital anti-inflammatory action on sepsis through activating autonomic nerve anti-inflammatory pathways, such as sympathoadrenal medullary pathway, but the mechanism remains unclear. This study aims to explore the optimum parameter of electroacupuncture (EA) stimulation in regulating sympathoadrenal medullary pathway and evaluate EA’s anti-inflammatory effect on sepsis. To explore the optimum parameter of EA at homosegmental acupoint on adrenal sympathetic activity, the left adrenal sympathetic nerve firing rate evoked by different intensities of single shock electrical stimulation (ES) at ST25 in healthy male Sprague-Dawley (SD) rats were evaluated by *in vivo* electrophysiological recording, and the levels of norepinephrine (NE) and its metabolites were also examined using mass spectrometry. To verify the role of EA at ST25 in sepsis, the rat was given intraperitoneal injection lipopolysaccharide to induce sepsis model, and survival rate, clinical score, and the level of interleukin (IL)-6, IL-1β, and IL-10 were evaluated after EA application. We observed that 3 mA is the optimal intensity on activating adrenal sympathetic nerve, which significantly elevated the level of NE in the peripheral blood. For LPS-treated rats, EA at the ST25 apparently increased the survival rate and improved the clinical score compared to the control group. Furthermore, 3 mA EA at ST25 significantly decreased pro-inflammatory cytokines IL-6 and IL-1β and upregulated anti-inflammatory cytokine IL-10 compared to the Lipopolysaccharide (LPS)-treated group. Overall, these data suggest that 3 mA is the optimal EA intensity at ST25 to activate the sympathoadrenal medullary pathway and exert an anti-inflammatory effect on sepsis.

**Highlights:**

- 3 mA single shock ES at ST25 drove adrenal sympathetic nerve reflexs.
- 3 mA EA at ST25 upregulated NE concentration in peripheral blood.
- EA at ST25 improved the survival rate in a rat of LPS-induced sepsis model.
- EA at the ST25 upregulated anti-inflammatory IL10 and downregulated pro-inflammatory IL1β and IL-6 in LPS-treated rats.

## 1. Introduction

Sepsis is described as life-threatening viscera dysfunction due to the excessive host response to inflammation (Singer et al. (2016). It is most frequently encountered in patients who have suffered severe trauma, infections, or even major surgical procedures. There are 30 million patients who suffer from sepsis worldwide each year, a condition with a 25-40% mortality rate in hospitals (Acuña-Castroviejo et al., 2017; Huang, Cai, & Su, 2019; Vincent et al., 2014). Several studies indicated that death resulting from Coronavirus 2019, the global pandemic disease, was associated with sepsis, which causes septic shock, multiple organ dysfunction syndromes (MODS), and acute respiratory failure distress syndrome(da Silva Ramos, de Freitas, & Machado, 2021; H. Li et al., 2020). Typically, antibiotics and steroids are the main methods used in treating sepsis, as microbial and viral infections primarily cause it. However, they were unsatisfactory because of severe side effects and sequelae. For example, long-term usage of antibiotics will destroy the microbiota and results in immunological dysregulation, ultimately leading to MODS, including the lungs, kidneys, and brain (Perner et al., 2016).

Recent studies showed that non-pharmacological approaches, especially acupuncture, positively affect sepsis (Liu et al., 2021; Liu et al., 2020; Song et al., 2012; Xie et al., 2020). Acupuncture has been widely used globally, and current progress revealed that the anti-inflammation action of acupuncture is predominately achieved by activating the autonomic nervous anti-inflammation pathways (sympathetic- and vagal-splenic pathway, vagal-adrenal medulla-dopamine pathway), rather than hypothalamic-pituitary-adrenal cortex axis (Ma, 2022; Pan, Fan, Chen, & Alemi, 2021). Moreover, increasing studies showed that the sympathetic nerve pathway contributes to neuroimmune regulation under pathological conditions (Scott-Solomon, Boehm, & Kuruvilla, 2021; Udit, Blake, & Chiu, 2022). The sympathoadrenal medullary axis engaged in the anti-inflammatory effect of acupuncture on inflammatory pain animal models (Arribas-Blázquez et al., 2020; Kim et al., 2008). Herein, we questioned whether the sympathoadrenal medullary axis involves in acupuncture-induced systemic antiinflammatory effect. Our previous studies observed segmental differences in the modulatory effects of acupuncture on target organs, with homosegmental acupoint stimulation preferentially evoking sympathetic nerves while heterosegmental acupoint stimulation mostly provoking the vagal nerve (X. Y. Gao et al., 2011; Y. Q. Li, Zhu, Rong, Ben, & Li, 2006; Su et al., 2013). Thereby, acupoint ST25, located at the same somatotopy as the adrenal gland, was chosen as the stimulating acupoint in the current study, exploring the optimum parameter of electroacupuncture (EA) stimulation for modulating adrenal sympathetic activity.

The adrenal glands are critical endocrine organs of the body to maintain physical homeostasis under physiological and pathological states by releasing medulla hormones and corticosteroids. The adrenal medulla is a significant source of catecholamines, which include norepinephrine (NE), epinephrine (E), and dopamine (DA). Numerous evidences demonstrated that NE acts as a neurotransmitter and is involved in the modulation of inflammation response in a concentration-dependent manner (Hasegawa et al., 2021; Lorton & Bellinger, 2015). A higher concentration of NE produces an anti-inflammatory action via binding β2-adrenoceptors (ADOB) expressed on immune cells, which subsequently leads to the release of antiinflammatory cytokines; in contrast, a lower concentration of NE exacerbates inflammation by binding α-adrenoceptors expressed on immune cells (Pongratz & Straub, 2014). Hence, we assumed that ES at ST25, a homosegemental acupoint to adrenal glands, can upregulate the peripheral concentration of NE via activating the sympathoadrenal medullary axis.

To address these questions, we firstly study the impact of different intensity ES at homosegmental acupoint ST25 on adrenal sympathetic activity in a physiological state, verifying that 3 mA ES at ST25 is the optimum intensity for switching on adrenal sympathetic activity, which can elevate the concentration of NE in peripheral blood. Ultimately, 3 mA EA at ST25 exerted an anti-inflammatory effect in a rat model of sepsis.

## 2. Experimental procedures

### 2.1 Animals

Healthy male SD rats (6-8 weeks old) weighing approximately 220 g were provided by the Experimental Center of China Academy of Military Medical Sciences. Rats were bred at the Institute of Acupuncture and Moxibustion, China Academy of Chinese Medical Sciences (CACMS), under the following conditions: air temperature 18-25°C, air humidity 40%-60%, 12h/12h alternating light-dark cycles, ad libitum water, and food intake for one week of adaptive feeding. Experiments were conducted in full compliance with the Guide for the Care and Use of Laboratory Animals of the National Institutes of Health. This protocol was approved by the Committee on Ethics of Animal Experiments of the Institute of Acupuncture and Moxibustion, CACMS.

### 2.2 Electrophysiological experiments

To assess the impact of EA at ST25 on adrenal sympathetic nerve activity in rat, somatic single shock electrical stimulation (ES) evoked adrenal sympathetic reflexes were recruited in the current study as previous described (Kimura, Sato, Sato, & Suzuki, 1996). Briefly, rats were anesthetized with an intraperitoneal injection of 50 mg/kg sodium pentobarbital (Sigma-Aldrich, St. Louis, MO, USA). An incision of approximately 1 cm in length was made along the midline, and the left abdominal muscles were bluntly dissected to expose the abdominal cavity. Next, the left adrenal sympathetic nerve, branches of the splanchnic nerve, were freed from surrounding connective tissues. After removed the peri lemma using fine forceps under microscope, adrenal sympathetic nerve was placed on a bipolar platinum hook electrode. The reference electrode was inserted into the subcutaneous tissue the incision site. The signal was amplified by amplifier (NL900D, Digitimer Limited, England) and collected through by an acquisition system with Spike 2 (CED1401, Cambridge Electronic Design, England). Filtering parameter: 300 Hz~1000 Hz, sampling frequency was: 50 kHz/s. For stimulating, a pair of needle electrodes were inserted subcutaneously into the acupoint ST25, which delivered the electrical stimuli with single square waves (parameters: pulse width: 1 ms, intensity: 1-4 mA, intervals of 10 min) from stimulator (STG4000, Warner Instruments, U.S.A) to evoke adrenal sympathetic reflex.

### 2.3 Rat sepsis model and clinical score

To prepare the Lipopolysaccharide (LPS)-induced sepsis model, rats were anesthetized by isoflurane and then given intraperitoneally injection of LPS (10 mg/kg). Clinical score: Rats were closely observed for dynamic changes at 1, 3, 6, 8, 24, 48, and 72 hours after LPS, injection and were scored according to Kadl et al (Kadl, Pontiller, Exner, & Leitinger, 2007). In details, the condition of conjunctivitis, stool consistency, hair coat, and activity upon moderate stimulation were assessed.

### 2.4 Electroacupuncture (EA) application

Rats were placed in a prone position on a heating pad and anesthetized with inhalation of 2% isoflurane. The bilateral ST25 (Tian shu) acupoints are located 5 mm lateral to the intersection between the upper 2/3 and the lower 1/3 of the line between the xiphoid process and the pubic symphysis upper border. The needles (0.25×25 mm, Suzhou Medical Appliance Factory, Suzhou, China) were inserted into the bilateral ST25 acupoints at a depth of ~3 mm. Rats were electrically stimulated for 20 min with 15 Hz in frequency, a pulse width of 1 ms, and 3 mA by a stimulator (STG4000, Warner Instruments, U.S.A) for three consecutive days.

### 2.5 Carotid artery blood collection

Rats were anesthesia by intraperitoneal injection with 50 mg/kg sodium pentobarbital (Sigma-Aldrich, St. Louis, MO, USA). Hair was removed from the neck’s midline. In the midline of the neck, a longitudinal incision was made approximately 1 cm long and the muscle was dissected bluntly using scissors to expose the thyroid gland. Carotid artery pulsation was observed between the sternocleidomastoid muscle and the thyroid cartilage of the larynx. The carotid artery blood was collected at the indicated time points, clotted at room temperature for 20 min, and centrifuged at 3000 g for 15 minutes at 4°C. The supernatant serum was collected and stored at −80°C before further experiment.

### 2.6 Mass spectrometry

Samples were thawed at 4°C for 30 minutes, and 200 μL serum or standard solution was transferred into a 1.5-mL centrifuge tube. Then, 200 μL internal standard solution was added, mixed by vortex, and centrifuged. The 350 μL supernatant was added to the activated solidphase extraction plate. Under positive pressure, impurities were eluted with 200 μL water and 200 μL 85% acetonitrile aqueous solution (containing 1% formic acid). Then, 40 μL 85% acetonitrile aqueous solution (containing 1% formic acid) was added to each well and treated with positive pressure to elute the target compound. The eluate was dried with nitrogen and reconstituted with a 1% formic acid aqueous solution. The receiver plate was placed on a thermostatic shaker and mixed for 1 min for mass spectrometry.

### 2.7 Enzyme-linked immunosorbent assay (ELISA)

ELISA was performed strictly according to the manufacturer’s instructions. Add wash buffer 350 μL/well, aspirate each well after holding 40 seconds, repeating the process two times for three washes. When loading the sample, 100 μl of the corresponding standard concentration was added in each of the 8standard wells, adding 100 μL of different concentrations of standard or sample in other wells, covered with the adhesive strip provided. Incubate for 2 hours at 37°C. Record the plate layout of standards and sample assay. Add 100 μL Working Biotin Conjugate Antibody (Rat IL-6 ELISA Kit, RK00020; Rat IL-1β ELISA Kit, RK00009; Rat IL-10 ELISA Kit, RK00050, Abclonal, China) in each well, cover with the new adhesive Sealer provided. Incubate for 1 hour at 37°C. Add 100 μL Working Streptavidin-HRP to each well, and cover with the new adhesive Sealer provided. Incubate for 0.5 hours at 37°C, plates were washed and dried, and 100 μL TMB Substrate to each well. Incubate for 15-20 minutes at 37°C. Protect from light; 50 μl Stop Solution was added to each well after 20 minutes of color development at 37°C. Plates were immediately put into the microplate reader, and optical density (OD) at 450/570nm was measured within 5 minutes. The average OD values for each standard protein concentration, quality control, sample, and other replicate wells were calculated, and IL-6 and IL-10 concentrations were calculated based on standard curves.

### 2.8 Statistical analysis

All data were subjected to the Shapiro-Wilk test to determine their normality. Only those data that passed the Shapiro-Wilk test have been represented in the graphs as mean SEM. Every figure legend explains how the methodology was used in each study. GraphPad Prism 8 (GraphPad Software, Inc., San Diego, CA, USA) was used for all analyses, and only *P*<0.05 was considered statistically significant.

## 3. Results

### 3.1 3 mA single shock ES at ST25 drove adrenal sympathetic nerve reflexs

To explore the intensity rule of ES at homosegmental acupoint ST25 in activating the sympathoadrenal medullary axis, somatic single shock ES evoked adrenal sympathetic nerve reflexes were recruited in the current study as previous described (Kimura et al., 1996). As depicted in Figure 1A-a, different intensity of signal shock ES (intensity: 1-4 mA, pulse width: 1 ms, intervals of 10 min) were applied to ST25 in healthy rats. The typical of adrenal sympathetic evoked by ES at ST25 is comprised A- and C-reflexes components and shown in Figure 1B-b. Generally, the A and C reflexes of sympathetic nerve evoked by somatic stimulation reflects the activation of the somatosympathetic reflex (Zhu, 2021). A-reflex has been related to spinal mechanism while C-reflex is associated with the spinal and supraspinal mechanism. Through calculating the frequencies of the adrenal sympathetic discharging induced by ES at ST25 acupoint, as well as the fold change of the discharge frequency relative to baseline, we observed that only the 3-mA ES intervention (frequency: 4.98±1.25 Hz, fold change to baseline: 181.2 ±40.44%) resulted in a significant elevation in discharge frequency compared to baseline (frequency:1.66±0.18 Hz) (Figure 1C-D, *P* < 0.01). Moreover, neither of low intensity in 1, 2 mA (frequency: 1 mA: 3.0±0.39 Hz, 2 mA: 2.79 ± 0.29 Hz, fold change to baseline: 1 mA: 92.64 ± 28.02%, 2 mA: 69.99 ± 7.556%) and high intensity in 4 mA ES (frequency: 3.0±0.3 Hz, fold change to baseline: 101.3 ± 45.37 %) altered adrenal sympathetic excitation compared to baseline (Figure 1C-D, *P* > 0.05). Together, these results suggested that 3 mA ES at ST25 is the optimal intensity for driving the adrenal sympathetic pathway. Hence, 3 mA was utilized as EA’s intensity in the next treatment study.

**Fig. 1.**
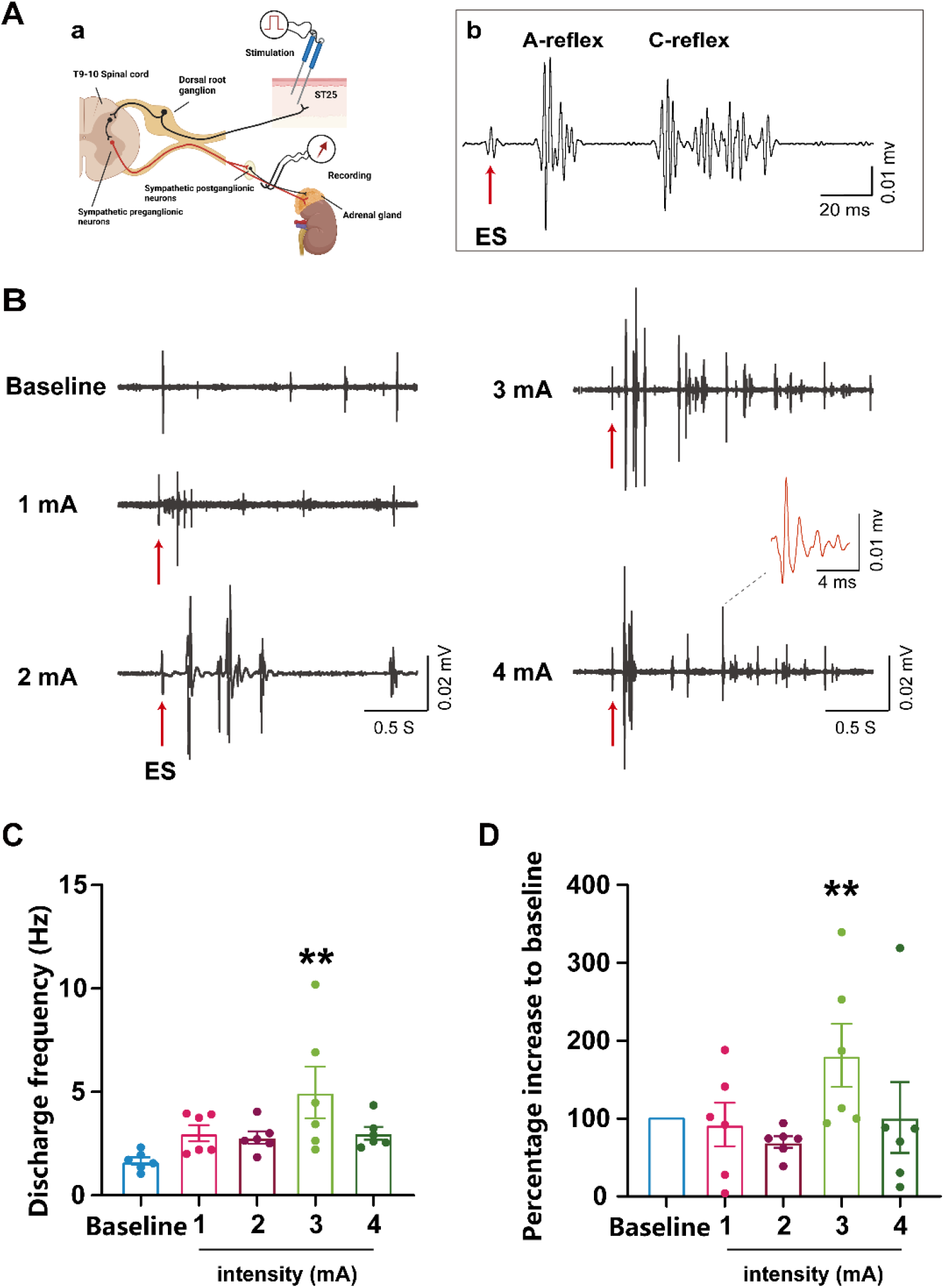
Different intensities ES at ST25 induced adrenal sympathetic nerve reflexes. A. (a) Schematic diagram of the adrenal sympathetic nerve reflexes induced by somatic single shock ES at ST25. A pair of needle electrodes were subcutaneously inserted into ST25 and used to stimulate while recording adrenal sympathetic nerve A- and C-reflexes with a bipolar platinum hook electrode. Created with BioRender.com. (b) Representative trace of ES at the ST25 evoked the adrenal sympathetic A- and C-reflexes. B. Representative trace of adrenal sympathetic A- and C-reflexes in response to 1-4 mA ES at ST25. C-D. Graphs showing the frequency of adrenal sympathetic discharge and the percentage change after in response to 1-4 mA ES at ST25. (n = 6/group, ** *P* < 0.01 vs. Baseline). One-way ANOVA with the least significant difference (LSD) multiple comparison tests.

### 3.2 3 mA EA at ST25 upregulated NE concentration in peripheral blood

Increasing evidence indicated that NE, one of adrenal medulla hormones and mainly neurotransmitter released by sympathetic postganglionic fibers or neurons, has been involved in the modulation of inflammation response in a concentration-dependent manner. Hence, to further explore whether peripheral NE was affected by 3 mA EA application, mass spectrometry was utilized to detect the concentration of NE and its metabolite, 3-methylnorepinephrine (3-MNE), in the peripheral blood after continuous 3 mA EA (15 Hz, 1ms pulse width, 20 min) at ST25. To avoid circadian oscillation of hormones,(Sharma & Farrar, 2020) we performed EA at 2 pm every day. As shown in Figure 2 A, 3 mA EA at ST25 significantly upregulated the NE levels in the peripheral blood in comparison with that in control rats, and we also observed a trend of increasing 3-MNE in the EA group. Hence, our results suggested that stimulating homosegmental acupoint ST25 by 3 mA EA can drive the release of peripheral NE from the sympathoadrenal medullary axis.

**Fig. 2.**
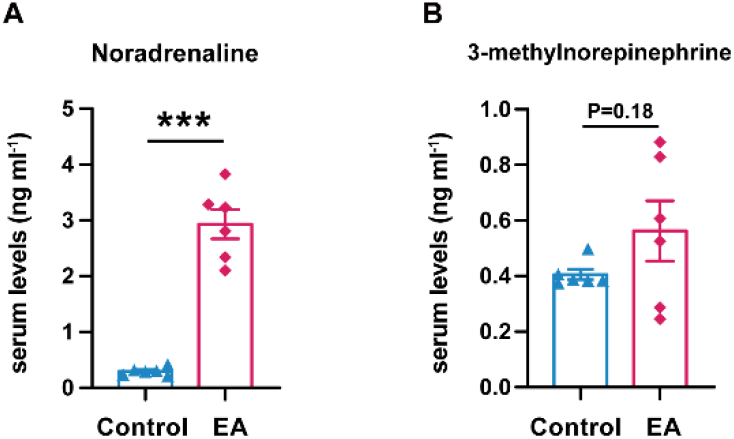
3 mA EA at the ST25 elevated NE (A) and its metabolite 3-methylnorepinephrine (B) in the peripheral blood. (n = 6/group, \*\*\**P* < 0.001 vs. Control). Independent t-test.

### 3.3 EA at ST25 improved the survival rate in a rat of LPS-induced sepsis model

Previous studies have suggested that the sympathoadrenal medullary axis are involved in the anti-inflammatory mechanisms of peripheral nerve stimulation (Willemze et al., 2018). But its’s role in acupuncture anti-inflammation effect remains exclusive. Given that increasing studies indicated that EA exhibited an anti-inflammatory effect on LPS-induced sepsis model, hence, we attempted to explore whether the sympathoadrenal medullary axis activated by EA at ST25 produced an anti-inflammation action in sepsis as well. In this part, 3 mA EA was applied to rats immediately after LPS injection and conducted for consecutive 3 days. As shown in Figure 3A, we observed that LPS administration resulted in most of rats in model group dead (survival rate: 16.67%, *P* < 0.001), whereas EA treatment dramatically enhanced the survival rate compared to model group (survival rate: 66.67%, *P* < 0.05). Meanwhile, the clinical scores of rats in either model or EA group were markedly decreased since 6 h after sepsis administration compared to that of control group (model: *P* < 0.001; ES: *P* < 0.05, Figure 3B); while EA treatment at ST25 greatly ameliorated the clinical symptoms of the rats, the clinical scores in EA group was significantly improved compared to model (*P* < 0.01). However, although LPS lead to the weight loss in model and EA group 2 and 3 days after sepsis establishment, there was no significantly change among these groups. Together, these data showed that EA treatment at ST25 produced a therapeutic effect on survival and clinical symptoms in sepsis rat model.

**Fig. 3.**
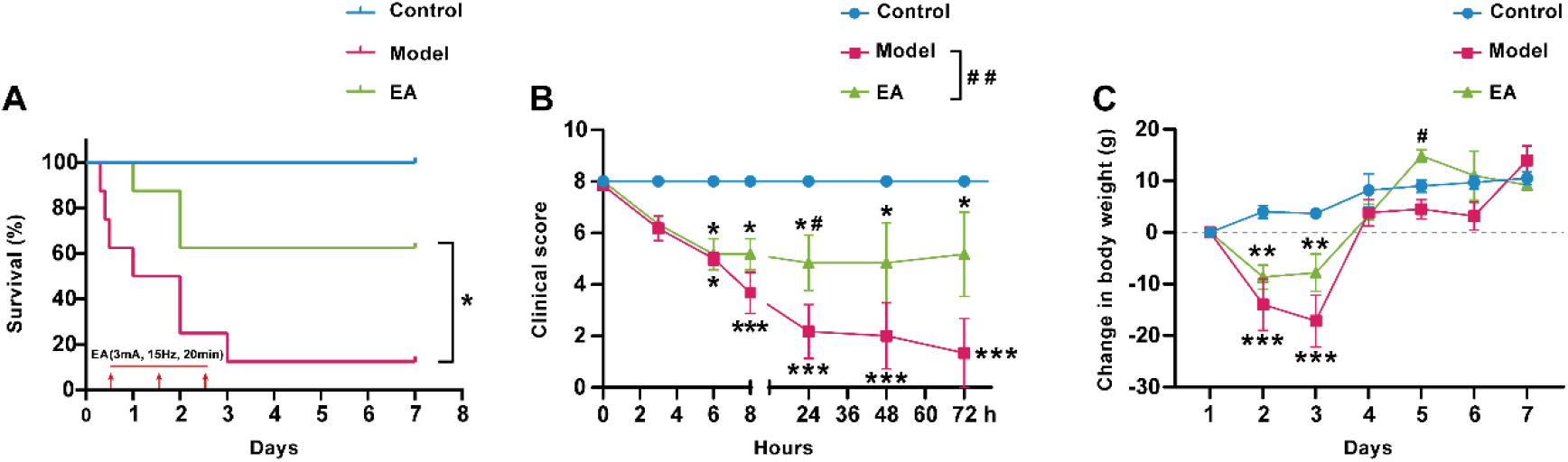
Effect of 3mA EA on survival rates, clinical score, and body weight in each group. A. 3mA EA at ST25 improved survival rates of rat after sepsis modeling. Arrows represent the time of EA treatment. (n = 8/group, * *P* < 0.05, vs. Model). Kaplan-Meier analysis. B. 3mA EA at ST25 improved clinical scores of rats after sepsis modeling. C. Comparison in body weight changes of rats in each group. (n = 8/group, * *P* < 0.05, ** *P* < 0.01, *** *P* < 0.001 vs. Control. # *P* < 0.05, ## *P* < 0.01, ### *P* < 0.001 vs. Model). Two-way ANOVA with Bonferroni test was to compare between groups.

### 3.4 EA at the ST25 upregulated anti-inflammatory IL10 and downregulated pro-inflammatory IL1β and IL-6 in LPS-treated rats

To explore the mechanism of EA treatment at ST25-mediated therapeutic effect on sepsis, we examined the change of the pro-inflammatory factors IL-6, IL-1β, and the anti-inflammatory factor IL-10 in the peripheral blood. As shown in Figures 4, LPS administration resulted in elevation of peripheral pro-inflammatory IL-6 and IL-1β, which were significantly higher than that in control (*P* < 0.001), indicating the LPS induced systematic inflammation. Notably, we observed EA treatment at ST25 not only decreased the level of peripheral pro-inflammatory IL-6 and IL-1β, but also altered the anti-inflammatory factor IL-10 level, which were significantly increased in rats received EA compared to model (*P* < 0.01). Collectively, these results suggested that 3 mA EA at ST25 produced a therapeutic effect on sepsis through decreasing pro-inflammatory factors and increasing anti-inflammatory factor.

**Fig. 4.**
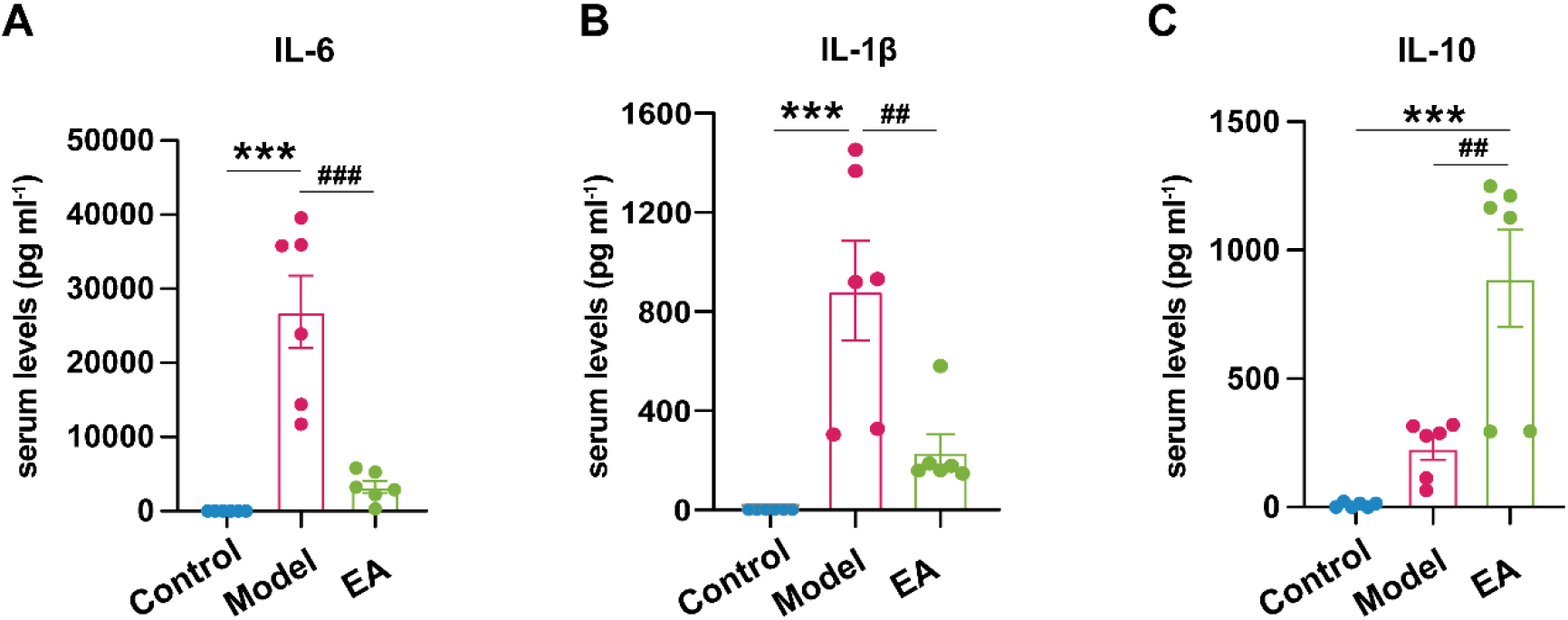
Levels of IL6 (A), IL-1β (B) and IL-10 (C) in the peripheral blood of rats in each group. (n = 6/group, *** *P* < 0.001vs. Control, ## *P* < 0.01, ### *P* < 0.001vs. Model). One-way ANOVA with the LSD multiple comparison tests.

## 4. Discussion

The major findings of this study were 3 mA of EA at ST25 served as the optimum stimulation intensity for triggering the sympathoadrenal medullary axis, which lead to increase of peripheral NE and then exert an anti-inflammatory effect in a rat model of sepsis. Moreover, 3 mA of EA at ST25 dramatically improved the survival rate and clinical score in sepsis modeling rats by altering the release of anti-inflammatory factors IL-10. These findings collectively demonstrated that 3 mA EA at ST25 can be used as a potential approach to modulated adrenal sympathetic activity and to treat inflammatory illnesses.

The adrenal glands are essential portion in the automatic nerve mediated anti-inflammatory pathway and contribute to the maintenance of the body’s homeostasis. The adrenal glands mainly consist of the cortex and the medulla, which the medulla is capable of releasing catecholamines, including NE, E, and DA. In contrast, the cortex mainly secretes mineralocorticoid, glucocorticoid, and gonadal hormones, which are involved in the endocrine regulation of the body. Neurodevelopmental evidence showed that the embryonic origin of the adrenal medulla is identical to that of sympathetic postganglionic neurons, which is functionally equivalent to sympathetic postganglionic neurons without axons and still is innervated by sympathetic preganglionic fibers (Scott-Solomon et al., 2021). Therefore, the adrenal medulla is often considered to belong to both the endocrine and autonomous nervous systems. Existing evidence shows that chromaffin cells in the adrenal medulla are directly innervated by the T9 segments of the preganglionic neurons. Kesse et al. found that the primary proportion of preganglionic fibers in the adrenal glands was innervated by T9, while T9 and T10 innervated a comparable proportion of postganglionic fibers (Kesse, Parker, & Coupland, 1988). We have previously observed a modulation pattern of “homotopic-acupoint” and “heterotopic-acupoint” that regulate visceral sympathetic and vagal activity (Y. Q. Li, Zhu, Rong, Ben, & Li, 2007). In detail, the acupoints that activate sympathetic nerve activity in the viscera are primarily located at the same dermatome as the viscera. In contrast, the acupoints that activate the vague nerveaare mainly located at the extremities relative to the distal segments of the viscera. In line with these neuromodulatory rules, our data also revealed that the acupoint ST25 (innervated by T10 spinal segment) (X. Gao et al., 2015), located at the same dermatome as the adrenal medulla, exhibited a better action in upregulating adrenal medulla hormones releasing via activating the adrenal sympathetic nerve (Fig1, 2).

Accumulating studies suggested that somatic stimulation exerted a neuromodulation action on sympathetic activity in a somatotopy-, intensity- and frequency-dependent manners (Cui et al., 2020; Kim et al., 2007; Liu et al., 2020; Takahashi & Imai, 2021). Sato et al. systematically studied the effects of different intensities of somatic stimulation on adrenal sympathetic activity and catecholamine concentration in the peripheral blood of rats. They showed that only noxious intensity stimulation on the abdominal skin significantly activated the adrenal sympathetic activity and increased the concentration of catecholamines in the peripheral blood (Sato, 1987). Through comparing the change of sympathetic reflex in response to different stimulation intensities, our data directly revealed that 3 mA ES at ST25 was the optimum intensity to trigger adrenal sympathetic activity; we did not observe 4 mA exert a better action on adrenal sympathetic activity. Previous studies have indicated that C-fibers, which contributed to the acupuncture therapeutic effect, can be activated by 3 mA of electrostimulation (Cui et al., 2022; Zhang et al., 2019; Zhu, Xu, Rong, Ben, & Gao, 2004). Furthermore, activated C fiber can drive the somato-sympathetic reflexes and enhance targeted visceral functions. Hence, the role of C fiber in these neuromodulatory effects is required to be investigated.

A recent study showed that the therapeutic effect of EA at ST25 on sepsis is disease-state-dependent. Liu et al. observed that 3 mA EA stimulation at ST25 1.5 h after sepsis modeling exacerbated inflammation reaction in mice, whereas 3 mA EA pretreatment at ST25 produced a significant anti-inflammatory effect via activating somato-spinal-splenic sympathetic pathway (Liu et al., 2020). Otherwise, we applied EA treatment to rat after sepsis modeling, which improved systematic inflammatory conditions. Hence, whether automatic nerve mediated anti-inflammatory pathways is different in rats and mice required to be further explored. Additionally, they and the other research groups reported that although high intensity (3 mA or 4 mA) EA applied at extremity acupoints (ST36 and LI4, whose location is heterosegment to adrenal gland), also can drive the spinal sympathetic reflex, the vagal nerve system primarily mediates the anti-inflammatory effect followed by EA application (Liu et al., 2021; Liu et al., 2020; Song et al., 2012). Concern the present study mainly focuses on regulating adrenal sympathetic activity and producing anti-inflammatory action through stimulating homosegmental acupoint of the adrenal gland. Hence, more studies are required to explore how to regulate adrenal function via manipulating hetersegmetal acupoint and exerting a better anti-inflammatory action and the underlying mechanism. Besides, although the vague-adrenal axis and the cholinergic anti-inflammatory pathway have been reported to mediate the anti-inflammatory effects of acupuncture at ST36, increasing evidence showed that the spleen, the cholinergic anti-inflammatory pathway’s central organ, is the cholinergic anti-inflammatory pathway closely related to the sympathetic nervous system (Elenkov, Haskó, Kovács, & Vizi, 1995; Katafuchi, Ichijo, Take, & Hori, 1993; Kees, Pongratz, Kees, Schölmerich, & Straub, 2003; Vida, Peña, Deitch, & Ulloa, 2011; Willemze et al., 2018). Therefore, exploring the role of the sympathetic nerve in the immune system and uncovering the neuromodulatory rules of acupuncture stimulation on adrenal sympathetic is beneficial to enhance the application of acupuncture on the treatment of sepsis and inflammatory illness.

Additionally, the current study showed that activating the sympathoadrenal medullary axis resulted in NE concentration upregulation in the peripheral blood. Interestingly, accumulating evidence has demonstrated that NE was involved in the inflammation response in a concentration-dependent manner. A high concentration of NE is inclined to raise antiinflammatory factor IL-10 released through binding ADOB expressed on immune cells, especially macrophages; in contrast, a low concentration of NE is apt to lead to the elevation of pro-inflammation tumor necrosis factor via binding with α-adrenoceptors on immune cells and exacerbate inflammation condition (Hasegawa et al., 2021; Pongratz & Straub, 2014). In line with our expectation, we observed that 3 mA EA treatment not only increased the survival rate, but it also decreased the pro-inflammatory factors IL-6 and IL1β and increased the level of the anti-inflammatory factor IL-10. Given that we have observed that 3 mA EA treatment resulted in a higher concentration of NE, thus, we assumed that the increased NE bonded with ADOB and then promoted the release of IL-10. However, how and where the increased NE impacts immune cells and leads to the anti-inflammatory factors release following 3 mA EA at ST25 required further exploration.

## Abbreviations

EA: electroacupuncture
SD: Sprague-Dawley
ES: electrical stimulation
NE: norepinephrine
IL-6: interleukin-6
IL-1β: interleukin-1 beta
IL-10: interleukin-10
LPS: Lipopolysaccharide
MODS: multiple organ dysfunction syndromes
E: epinephrine
DA: dopamine
ADOB: β2-adrenoceptors
ELISA: Enzyme-linked immunosorbent assay
3-MNE: 3-methylnorepinephrine
LSD: least significant difference

## Acknowledgements

We are grateful to Dr. Shufeng Li of Beijing Hexin Technology Co., Ltd. for technical assistance. We would like to thank Editage (www.editage.cn) for English language editing.

## Author contributions

B.Z., X. C., X. G. designed and directed the project. Z. Z., H. X., Q. Z., Y. Z., and D. Z. performed most of the experiments. Z. Z. and K.L. analyzed data. Z. Z. and X.C. was involved in writing the first draft manuscript. B.Z. and X. C. wrote the final manuscript

## Data availability

The data will be made available upon reasonable request.

